# Informational masking vs. crowding — A mid-level trade-off between auditory and visual processing

**DOI:** 10.1101/2021.04.21.440826

**Authors:** Min Zhang, Rachel N Denison, Denis G Pelli, Thuy Tien C Le, Antje Ihlefeld

## Abstract

In noisy or cluttered environments, sensory cortical mechanisms help combine auditory or visual features into perceived objects. Knowing that individuals vary greatly in their ability to suppress unwanted sensory information, and knowing that the sizes of auditory and visual cortical regions are correlated, we wondered whether there might be a corresponding relation between an individual’s ability to suppress auditory vs. visual interference. In *auditory masking*, background sound makes spoken words unrecognizable. When masking arises due to interference at central auditory processing stages, beyond the cochlea, it is called *informational* masking (IM). A strikingly similar phenomenon in vision, called *visual crowding*, occurs when nearby clutter makes a target object unrecognizable, despite being resolved at the retina. We here compare susceptibilities to auditory IM and visual crowding in the same participants. Surprisingly, across participants, we find a negative correlation (*R* = −0.7) between IM susceptibility and crowding susceptibility: Participants who have low susceptibility to IM tend to have high susceptibility to crowding, and vice versa. This reveals a mid-level trade-off between auditory and visual processing.

## Introduction

Hearing-impaired individuals frequently do not understand spoken words when background sound interferes. Often this occurs because the same neurons in the auditory nerve respond to both the target of interest and the background sound, swamping target information. However, many people, including those with “normal” audiological thresholds, can have trouble understanding speech even when background sound energy flanks, rather than overlaps, a target’s spectrum, a situation in which cochlear models predict no interference (***Goldsworthy and Greenberg, 2004***). This second difficulty with background sound is often attributed to an incompletely understood central auditory phenomenon called *informational masking* (*IM*); ***Kidd and Colburn*** (***2017***)). IM occurs when target and masker resemble each other or when the listener is uncertain about their perceptual properties (***Brungart, 2001***; ***Lutfi et al., 2013***). To characterize the mechanism underlying IM, we here ask whether and how IM relates to a phenomenologically similar effect in vision: crowding ***(Pelli et al., 2001)***. In crowding, clutter in the visual scene prevents object recognition. Using comparable crowding and IM tasks, we are surprised to find that crowding susceptibility correlates negatively with IM susceptibility, revealing a mid-level trade-off between auditory and visual processing.

As perceptual phenomena, IM and crowding are strikingly similar. Both are cortical processes by which target-like maskers can prevent target identification (***Scott et al., 2004**, **2006**, **2009***; ***Gutschalk et al., 2008***; ***Pelli, 2008***; ***Millin et al., 2014***). Both are refractory to learning in neurotypical individuals (***Neff et al., 1993***; ***Hussain et al., 2012***). In crowding, nearby clutter is perceived as part of the visual target (Figure 1A). Analogously, in IM, flanking sound, with little energy inside the target’s critical band, hampers individuation of target and flankers (Figure 1B). Crowding and IM are stronger when target and masker are perceptually similar. Both phenomena occur even when the masker is much fainter than the target, or when the target is presented to one eye/ear and the masker to the other eye/ear (***Kidd and Colburn, 2017**; **Ihlefeld and Shinn-Cunningham, 2008***; ***Pelli et al., 2001***; ***Gallun et al., 2007***; ***Flom et al., 1963***). In vision, clutter crowds a target when the target and clutter are less than a specific distance apart, called the crowding distance (or critical spacing). Crowding distance predicts an individual’s crowding susceptibility over a broad range of stimuli ***(Pelli et al.,2001)***. Given these functional similarities, we wondered whether IM and crowding rely on similar mechanisms.

**Figure 1.**
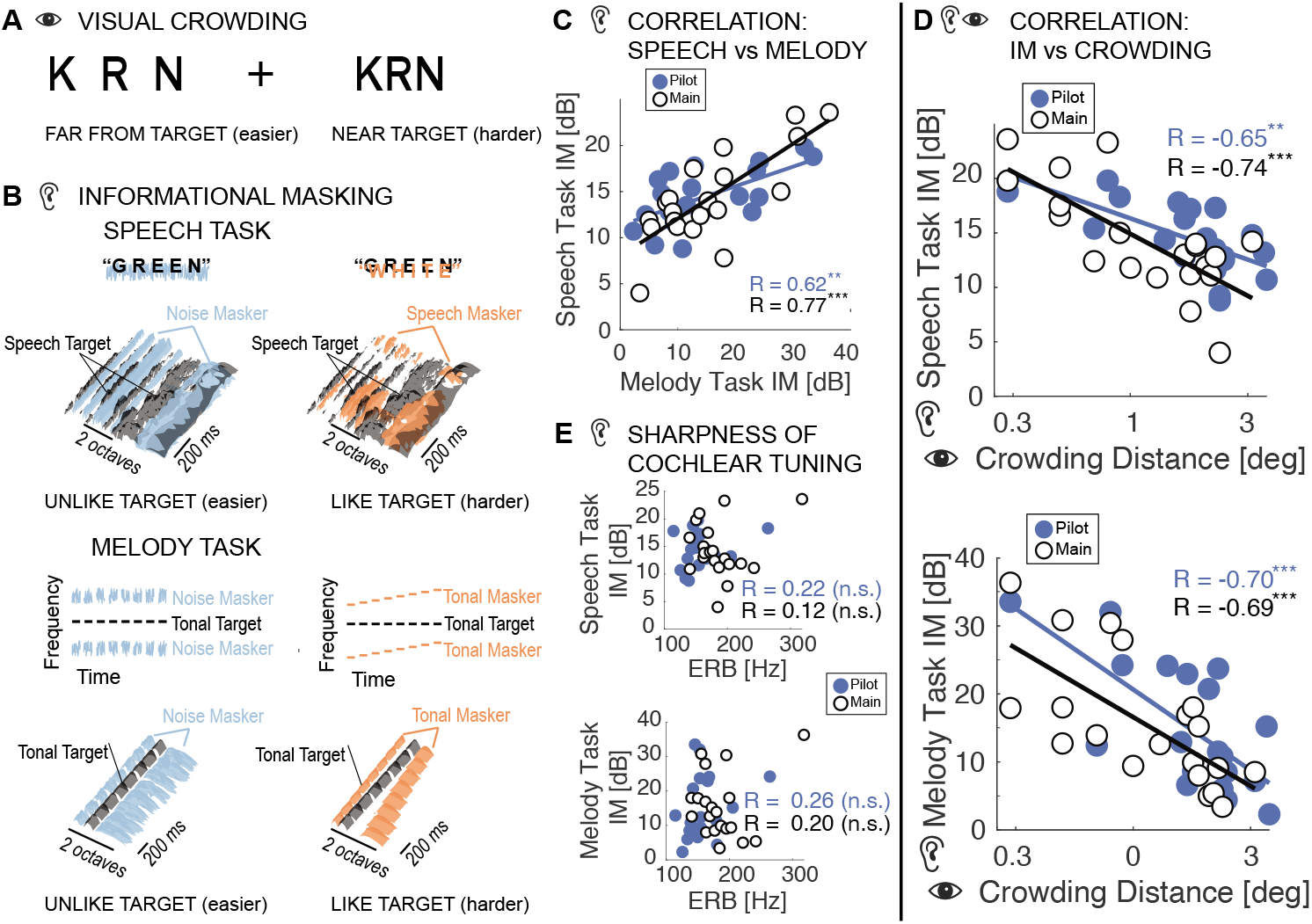
In all graphs: Susceptibility increases with increasing IM and crowding distance. Blue and white symbols represent the pilot and main experiments, respectively. (**A**) To experience crowding, fixate the cross. While fixating the cross, try to identify the middle letter in each triplet, ignoring the outer letters. When they are closer, the target is harder to identify. Without the outer letters, the two targets, left and right, would be equally visible. (**B**) Schematics and spectrograms of the sounds we used to study informational masking (IM). The noise masker covered the same spectral range as the speech masker in the speech task or the tonal masker in the melody task, such that masking in the cochlea should be greater or equal in the noise vs. speech or tonal masker conditions; any excess masking in the speech or tonal masker conditions is post-cochlear, i.e. IM. We measure the participants accuracy in identifying the target sound (denoted in black) as a function of the Target-to-Masker broadband energy ratio (TMR). Target identification is easier when the masker is unlike the target (left spectra). Therefore, threshold TMR for target-like masking is larger than that for noise masking. For each participant, the differences in threshold TMRs measure the individual’s IM susceptibility. (**C**) Participants who are IM-susceptible in the speech task also tend to be IM-susceptible in the melody task. (**D**) However, IM susceptibility in the speech task is anti-correlated with the participant’s crowding susceptibility. Similarly, participants who are less crowding-susceptible tend to be more IM-susceptible in the melody task. (**E**) Equivalent Rectangular Bandwidths (ERBs) at the target center frequency of 1000 Hz (estimated from noise-masking thresholds in unlike-target conditions of the melody task) confirm that participants had “normal” sharpness of peripheral tuning, with individual ERBs that were smaller than the smallest tested notch width. (Note that a 0.3 octave notch width corresponds to 208 Hz at 1000 Hz. One participant’s ERB exceeded 208 Hz, but removing this participant does not affect the conclusions.) ERBs are a poor predictor of IM susceptibility in the speech task. Similarly, ERBs and IM susceptibility in the melody task are uncorrelated.

## Results

We measured behavioral performance in one crowding and two IM tasks in the same 20 normal-hearing young participants. To measure crowding distance, participants identified a peripheral target letter between two flanking letters (Figure 1A). To quantify IM, we used a speech and a non-speech task. In both tasks, we separately tested maskers that were either target-like or noise, which is not target-like, and varied the *target-to-masker broadband energy ratio* (*TMR*). IM susceptibility is the difference between threshold TMRs in target-like background sound vs noise. In the “speech task,” participants identified target words constructed from narrow spectral bands that were masked by either speech (target-like) or noise. In the “melody task”, participants reported whether a target sequence of eight constant frequency tones was present while spectrally flanked by masking sequences of either target-like tones or noise bursts (Figure 1B).

We initially ran this experiment as a pilot study with 20 participants. Prompted by the unexpected findings, we re-ran it as the “main” experiment with a new group of 20 participants and minor modifications (see Appendix). Results of the pilot and main experiments are similar (compare blue vs white symbols in Figure 1C-E). IM susceptibility in the melody task predicts IM susceptibility in the speech task (pilot: ***R*** = 0.62, *p* = 0.003; main: ***R*** = 0.77, *p* < 0.001, Figure 1C), suggesting that IM susceptibility generalizes across speech and non-speech stimuli. Moreover, across participants, crowding susceptibility (i.e. crowding distance) and IM susceptibility are correlated, but, unexpectedly, the correlation is negative. In this way, crowding is inversely related to IM for both the speech task (pilot: ***R*** = −0.65, *p* = 0.002; main: ***R*** = −0.74, *p* < 0.001) and the melody task (pilot: ***R*** = −0.70, *p* < 0.001; main: ***R*** = −0.69, *p* < 0.001, Figure 1D). Equivalent rectangular bandwidth (ERB), estimated from noise masking thresholds, does not correlate with IM susceptibility (Speech: pilot: ***R*** = 0.22, *p* = 0.4; main: ***R*** = 0.12, *p* = 0.6; Melody: pilot: ***R*** = 0.26, *p* = 0.261; main: ***R*** = 0.20, *p* = 0.399, Figure 1E), confirming that an individual’s sharpness of cochlear tuning does not predict their IM susceptibility (***Oxenham et al., 2003***).

## Discussion

These results show that IM and crowding, two remarkably similar mid-level sensory processes, are related, but in an inverse fashion. The neural origin of this mid-level trade-off between auditory and visual processing is unclear. Shared mechanisms that induce positive correlations — including generic factors like developmental deprivation (***Hoddinott et al., 2013***), domain-general selective attention (***Shinn-Cunningham, 2008***; ***Clayton et al., 2016***), motivation, effort, or vigilance (***Pichora-Fuller et al., 2016***) cannot explain this effect. Assuming that crowding distance (in mm) is conserved in cortex, the sizes of underlying visual cortical areas should be reciprocally proportional to crowding distance (in deg; ***Pelli*** (***2008***)). Crowding distance correlates with size of hV4, but not V1 (***Kurzawaski et al., 2021***). The correlation between sizes of A1 and hV4 is unknown, but primary visual and auditory cortices tend to covary in size (***Song et al., 2011***). Visual areas V1, V2, V3, and hV4 are in posterior cortex, whereas auditory cortex is more anterior. Patients with posterior cortical atrophy are more susceptible to visual crowding and — surprisingly — less able to perceptually segregate auditory scenes (***Hardy et al., 2020***), unlike our negative correlation in neurotypical participants. Finally, we wonder if years of prolonged visual or auditory attention might reduce crowding or IM, respectively. People who spend more time looking may listen less, and vice versa. Further work is needed to discover how an individual’s ability to recognize a target in clutter develops in each sensory modality.

## Methods and Materials

### Participants

A total of 40 participants (ages 19 to 25) took part in the study, 20 per experiment (8 females in the main experiment, and 10 females in the pilot). All participants had normal audiometric pure-tone detection thresholds as assessed through standard audiometric testing at all octave frequencies from 250 Hz to 8 kHz. At each tested frequency, tone detection thresholds did not differ by more than 10 dB across ears, and all thresholds were 20 dB HL or better. All participants self-reported that they had never learned to play an instrument and never sung in a vocal ensemble. All participants gave written informed consent to participate in the study. All testing was approved by the Institutional Review Board of the New Jersey Institute of Technology.

### Experimental setup

Throughout testing, participants were seated inside a single-walled sound-attenuating chamber (International Acoustic Company, Inc.) with a quiet background sound level of less than 13 dBA. Acoustic stimuli were generated in Matlab (Release R2016a, The Mathworks, Inc., Natick, MA, USA), D/A converted with a sound card (Emotiva Stealth DC-1; 32 bit resolution, 44.1 kHz sampling frequency, Emotiva Audio Corporation, Franklin, TN, USA) and presented over insert earphones (ER-2, Etymotic Research Company Elk Grove Village, IL, USA). The acoustic setup was calibrated with a 2-cc coupler, 1/2” pressure-field microphone, and a sound level meter (2250-G4, Brüel and Kjær, Nærum, Denmark). Visual stimuli were delivered via a 23-inch monitor with 1920 x 1080 resolution. Prior to visual testing, the experimenter positioned the monitor such that it was centered 50 cm away from the center of the participant’s nose. Using this setup, three task were administered in an order that was counterbalanced across participants.

### Visual Crowding Task

Crowding distance was assessed with a target identification paradigm (***Pelli et al., 2016***), illustrated in Figure 2A. Participants were instructed to fixate their gaze at the center of a cross hair displayed on the monitor in front of them. A target letter was shown between two flanking letters. The three letters were placed in a row and randomly chosen without replacement from nine possible letters <D, H, K, N, O, R, S, V, Z>. Target and flankers appeared together to either the right or left of the cross hair for 0.5 seconds, then disappeared. Participants were tasked to read out loud the middle letter while continuing to fixate on the cross hair. An experimenter recorded the participant’s response on each trial. Using the QUEST method (***Watson and Pelli, 1983***), the distance between target and flanker letters was adaptively varied in two blocks of 20 trials, aiming for 70% correct and assuming a Weibull psychometric function. Each track started with an initial guess for distance from the center of the middle letter based on neurotypical critical spacing (***Song et al., 2014***). The crowding distance measured in the first track was a practice run. If estimated crowding distances between first and second track differed by more than 0.2 degrees, a third track of 20 trials was administered. The crowding distance on the final track was recorded as the participant’s threshold.

**Figure 2.**
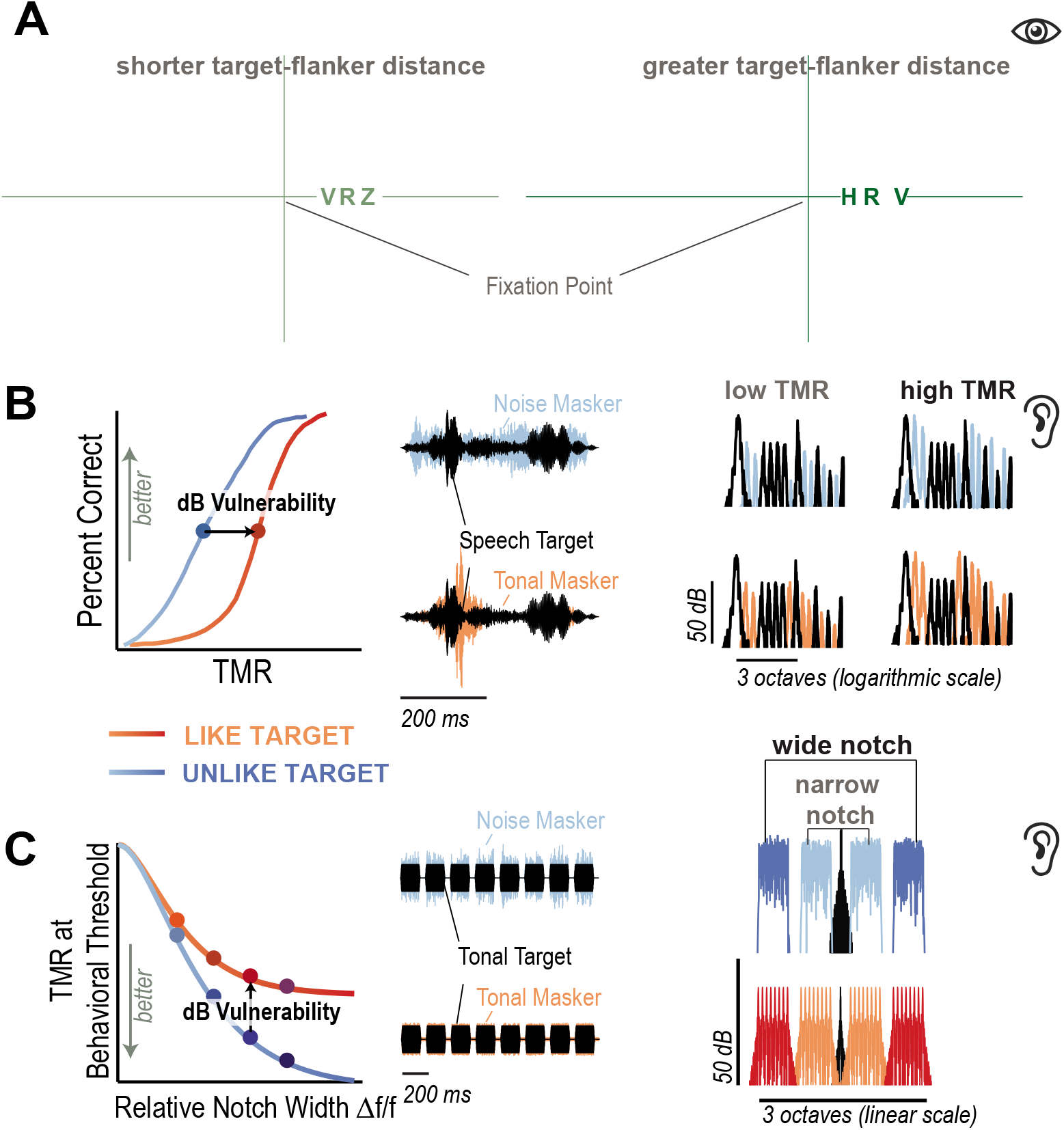
**(A)**In the visual crowding task, participants fixated the cross hair and called out the target letter in the center (here: *R*), while two flankers were cluttering it. Crowding distance was measured by adaptively varying the distance between the target and flankers. In the example here, the target and flanking letters are shown to the right side of the fixation cross hair. In the main experiment, the letters would randomly appear to the left or the right of the cross hair, whereas in the pilot experiment they only appeared to the right. **(B)** In the speech task, susceptibility to IM was measured by subtracting the TMR where participants correctly identified 50% of target words in the presence of background speech vs. background noise. **(C)** Analogously, in the melody task, susceptibility to IM was assessed as the difference in TMRs between target detection with melody maskers vs. noise maskers, and across different notch widths.

### Speech Task

IM susceptibility was measured in a speech task (Figure 2B), by subtracting the TMR at 50% correct speech identification threshold with a speech masker from the threshold TMR with a noise masker (***Arbogast et al., 2002***). Speech identification was assessed using the Coordinate Response Measure matrix task (***Bolia et al., 2000***). This matrix task uses sentences of the following fixed structure: ‘Ready [callsign] go to [color] [number] now.’ During testing, a target and a masker sentence were simultaneously presented to the left ear only, constrained to differ from each other in terms of call-sign, color and number keywords (***Kidd Jr et al., 2008***). Target sentences always had the callsign ‘Baron.’ There were four color keywords <‘red’,’blue’,’white’,’green’> and seven possible numbers <‘one’,’two’,’three’,’four’,’fíve’,’six’,’eight’> (excluding the number’seven’ because it has two syllables. Participants were instructed to answer the question: Where did Baron go?’ by pressing the corresponding color and number buttons on a touch screen response interface. A trial was counted as correct if the participant correctly reported the target color and the target number, resulting in a chance performance of 4% 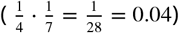.

To vocode the utterances, raw speech recordings were normalized in root mean square (RMS) value and filtered into 16 sharply tuned adjacent frequency bands using time reversal filtering, resulting in no appreciable phase shift. Each resulting band covered 0.37 mm along the cochlea between 3-dB down-points according to Greenwood’s function (***Greenwood, 1990***), or approximately 1/10th octave bandwidth, and had a 72 dB/octave frequency roll-off, with center frequencies ranging from 300 to 10,000 Hz. In each narrow speech band, the temporal envelope of that band was then extracted using the Hilbert transform and multiplied by uniformly distributed white noise carriers. To remove the side bands, the resulting amplitude-modulated noises were processed by the same sharply tuned filters that were used in the initial processing stage. Depending on the experimental condition, a subset of these sixteen bands was then added, generating intelligible, spectrally sparse, vocoded speech. Stimuli were generated from utterances by two different male talkers, one for the target and a different talker for the speech masker.

To generate noise maskers that matched the spectrum of the vocoded speech, all processing steps were the same as in the vocoding described above, with one exception. Instead of using the Hilbert envelope to amplitude-modulate the noise carriers, here, the noise carriers were gated on and off with 10 ms cosine-squared ramps that had the same RMS as the Hilbert envelope of the corresponding speech token in that band. A subset of the resulting 16 narrowband pulsed noise sequences was added to generate low-IM noise maskers.

On each trial, nine randomly chosen bands were added to create the target. The masker was comprised of the remaining seven bands and either consisted of vocoded utterances from the same corpus, recorded by a different male talker (target-like) or of noise tokens with similar long-term spectral energy as the vocoded utterances (target-unlike). The center and right panels of Figure 2B shows a representative temporal and spectral energy profile for a mixture of target (black) and speech (purple) or noise (brown) maskers.

The masker was presented at a fixed level of 55 dB SPL. The target level varied randomly from trial from 35 to 75 dB SPL, with a 10 dB step size, resulting in five broadband TMRs from −20 dB to 20 dB. For familiarization with the vocoded speech task and to ensure that the vocoded speech stimuli were indeed intelligible, participants were initially tested in 20 trials on nine-band target speech at 35 dB SPL, without masking. Target bands varied randomly from trial to trial. All participants reached at least 90% accuracy during this testing in quiet.

Next, participants were tested in five blocks of 40 trials while target-unlike noise or target-like speech interfered in the background. Thus, each specific combination of the five different TMRs and two masker types was presented 20 times (5 TMRs * 2 masker types *20 trials = 5 blocks *40 trials = 200 trials total). TMR and masker type varied randomly from trial to trial such that all combinations ofTMR masker type were presented in random order once before all of them were repeated in a different random order.

To estimate the TMR at the 50% correct threshold for each participant, percent correct scores as a function of TMR were fitted with Weibull-distributed psychometric functions, by using the `psignifit` package (***Wichmann and Hill, 2001***). IM susceptibility was computed as the difference in TMR at 50% correct between noise vs. speech masking.

### Melody Task

IM susceptibility was also assessed using two-up-one-down adaptive tracking with a non-speech task, by contrasting 70.7% correct thresholds for detecting eight-tone-burst targets across two notched-masker conditions: a noise vs. a melody masker (***Levitt, 1971***), illustrated in Figure 2C. Target and masker were presented to the left ear only. The target consisted of eight pure tones at a fixed frequency of 1000 Hz. Each tone was 150 ms long (including 10 ms cosine-squared ramps, random phase), with 75 ms gaps between consecutive tones. The target intensity was varied adaptively.

Using a classic paradigm for estimating ERB, in the noise masker condition, two 600-Hz-wide narrow bands of noise were placed symmetrically around the target frequency, creating a symmetrical notch in logarithmic frequency (***Patterson, 1976***). The notch width was one of the following: <0.3, 0.5,1,1.5> octaves. Noise tokens with very steep spectral slopes of over 400 dB/octave were constructed by generating uniformly distributed white noise, transforming it via Fast Fourier Transform and setting the notch frequencies in the spectrum to 0, before transforming the signal via the real portion of the inverse Fast Fourier Transform back into the time domain.

The melody masker condition was designed to closely match the spectral profile of the noise masker. Two eight-tone melodies, each carrying eight possible frequencies that were spaced linearly within 600-Hz-wide bands, flanked the target. One of these melodies was played above, the other below the target frequency, positioned symmetrically around the target frequency along a logarithmic frequency axis. The maskers were chosen from four possible melodies <up, down, up-down, down-up>. Those patterns indicated how the frequency changed for eight pure tones that formed the sequence, for instance ‘up’ means that each pure tone increased in frequency compared to the previous one in the sequence. The phase of each tone was independently and randomly drawn for each tone, resulting in phases that generally differed across all tones in the target-flanker mixture.

Maskers were played at a fixed spectrum level of 40 dB SPL, equivalent to a broadband level of 68 dB SPL (total level =40 + 10 · *log*_10_(300) + 10 · *log*_10_(2) = 68). To protect against the possibility of distortion products as a possible task cue in the melody condition, a low-intensity broadband white noise masker was continuously played in the background during both the noise and the melody masker condition at 15 dB SPL.

Under both masker conditions, participants performed a two-alternative forced-choice target detection task, responding with ‘yes’ or ‘no’ to indicate whether they heard the target. At the beginning of each adaptive track, the target intensity started at 70 dB SPL. The target intensity was initially decreased by 10 dB for every two consecutive correct answers and increase by 10 dB for every incorrect answer. After every two reversals in the adpative tracks, the step size was halved. Participants completed 12 reversals. Threshold was the average target intensity across the final 12 responses. IM susceptibility was calculated by subtracting thresholds between noise and melody masker at one octave separation.

### ERB

To estimate each participant’s ERB, noise masked thresholds from the melody task, denoted as *W*, were minimum-least-square fitted to rounded exponential (roex) functions (command `lsqcurvefit` in Matlab). The roex functions were defined as

*W*(*g*) = (1 − *r*)(1 +*pg*)*e^−pg^* + *r*, where *p* determined the steepness of the roex function’s passband, *r* shaped the stopband, and *g* denoted the distance between the target frequency *f_T_* and the corner frequencies of the masker notch with 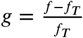 (***Patterson et al., 1982***; ***Oxenham et al., 2003***).

### Pilot Experiment

The methods used for the pilot experiment were similar to those of the main experiments except for two differences. First, unlike in the main experiments, the tasks in the pilot experiment were not counterbalanced across participants and administered in fixed order instead. Specifically, during pilot testing, participants first completed the crowding task, followed by the IM speech task, followed by the IM melody task. The second difference across the experiments is that in the crowding task during piloting, participants were only tested to the right side of the cross hair. Whereas during the main experiments, target and masker randomly alternated between presented to the left vs. the right side of the cross hair.

### Statistical Analyis

Statistical analyses were performed using linear regression via the command `fitlm` in Matlab 2019b, and R and adjusted R-squared from these fits are reported. Multiple comparisons were adjusted with Bonferroni correction.

## Acknowledgments

USA NIH grant R01DC019126 to Ihlefeld and R01EY027964 to Pelli funded this work. The authors thank Jonathan Winawer, Jan Kurzawski, Najib Majaj, Keir Yong, Jamie Radner, and Jasmine Kwasa for commenting on an earlier version of this manuscript.

## Appendix 1 Melody task

The majority of participants have ERBs at or below 208 Hz (Appendix 1 Figure 1A), with no appreciable differences in the sharpness of cochlear tuning between participant groups in the main vs. pilot experiment. As expected, thresholds were much less variable across listeners in the the noise masker condition as compared to the melody condition, in both the main and pilot experiment (compare the spread in the density plots in the top vs. bottom panels of Appendix 1 Figure 1B). Considering the high across-participant variability in susceptibility to IM, we next used bootstrapping to estimate the confidence intervals the correlation between crowding distance and susceptibility to non-speech IM, as a function of notch width (Figure Appendix1B). Adjusted correlation coefficients roughly increased with increasing notch width and were most consistent across the main vs. pilot experiments at the 1-octave notch width.

**Appendix 1 Figure 1.**
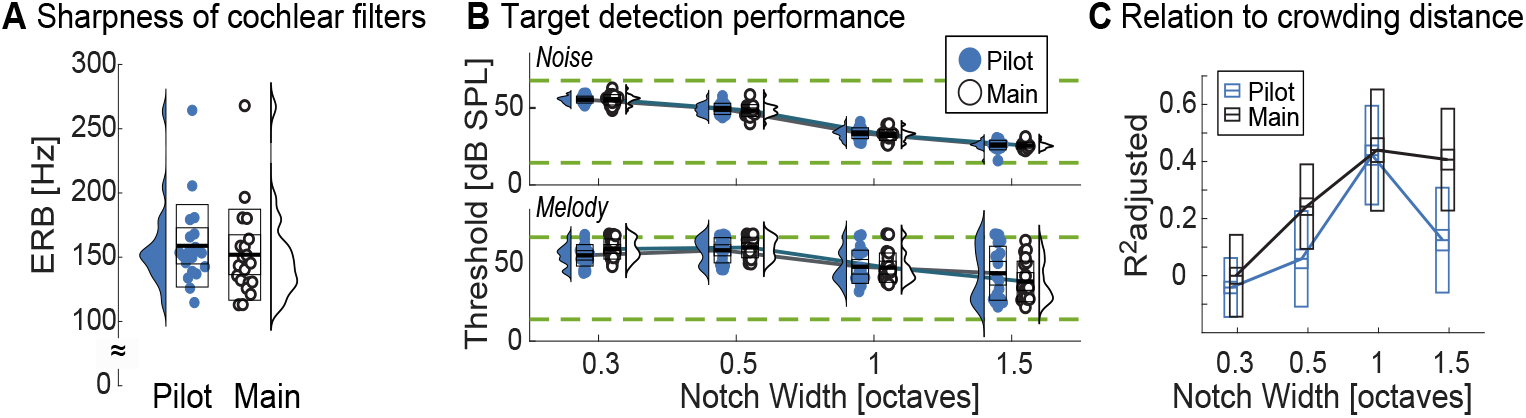
**(A)** The range of ERBs is comparable across both experiments. **(B)** Individual variability is much higher in target-like masking than noise across all tested ROIs in experiment 1. For all participants, target detection thresholds generally fall between the broadband level of the notched masker (68 dB SPL) and broadband masker (15 dB SPL), shown by green dashed lines. In the noise masker configuration, target detection thresholds decrease with increasing notch width for all participants (top), but only for some participants in the melody masker configuration (bottom). Note that in both the main and the pilot experiment, the densities in the melody task are bimodal, with one mode close to 68 dB SPL throughout, and the other mode decreasing with increasing notch width. **(C)** The adjusted correlation coefficient between visual crowding distance and IM susceptibility reveals coarse tuning. Results between the two experiments are congruent at 0.3 and 1 octave notch widths, but appear to diverge at 0.5 and 1.5 octave notch widths. Test-retest variability of ***R***^2^, estimated via bootstrapping that sampled 10 out of 20 participants without replacement 100 times, show that, indeed, crowding distance is robustly correlated with IM susceptibility at 1 octave separation, in both experiments. At other notch widths, the relationship is less pronounced.

Visual inspection of the density functions in Appendix 1 Figure 1B hints that the distribution of melody masking thresholds was bimodal, gradually widening with increasing octave separation. The mean of the lower mode, corresponding to participants who were more resilient to masking, decreased with increasing notch width. The mean of the other mode remained roughly constant as a function of notch width and close to the broadband level of the masker, indicating that the more poorly performing participants chose a strategy to listen for the louder source as opposed to relying on target pitch. In these poorly performing listeners, thresholds did not monotonically improve with increasing notch width. Perhaps as a result, roex functions used to estimate ERB under noise masking did not provide appropriate fits of the data under melody masking.

While we did not originally anticipate this result, in general, approximately a third of normal-hearing listeners have difficulty discerning pitch, and can, for instance, not reliably distinguish between major and minor triads in musical chords, even when given trial-by-trial correct response feedback (***Chubb et al., 2013***; ***Mednicoff et al., 2018***; ***Graves and Oxenham, 2019***). Note that we here tested the IM melody task at 0.3, 0.5,1 and 1.5 octave notch width, resulting in center frequencies of the lower and upper flanker bands that were related by factors of 1.231, 1.414, 2.000 and 2.828. Those numbers were originally chosen to cover the range of notch widths that typically result in ERB estimates (***Patterson et al., 1982***). However, they had the unintended effect that the constituent flanker frequencies were not perfectly harmonically related, and therefore potentially unfused at 0.3, 0.5 and 1.5 octaves, whereas flanker frequencies were harmonically related at the 1 octave notch. In summary, IM susceptibility in the melody task at these other three notch widths is more weakly correlated or even uncorrelated with crowding distance (as well as IM susceptibility to speech), showing that harmonicity affects IM in this paradigm. Moreover, domain-general selective attention (***Clayton et al., 2016***) or systemic developmental deprivation (***Hoddinott et al., 2013***) cannot account for this ***R***^2^-tuning nor for the inverse association between IM and crowding in the main experiment.

